# A Parallel Implementation of the Finite State Projection Algorithm for the Solution of the Chemical Master Equation

**DOI:** 10.1101/2020.06.30.180273

**Authors:** Huy Vo, Brian Munsky

## Abstract

Stochastic reaction networks are a popular modeling framework for biochemical processes that treat the molecular copy numbers within a single cell as a continuous time Markov chain, whose forward Chapman-Kolmogorov equation is known in biochemistry literature as the chemical master equation (CME). The solution of the CME contains extremely useful information that can be compared to experimental data in order to improve the quantitative understanding of biochemical reaction networks within the cell. However, this solution is costly to compute as it requires integrating an enormous system of differential equations that grows exponentially with the number of chemical species. To address this issue, we introduce a novel multiple-sinks Finite State Projection algorithm that approximates the CME with an adaptive sequence of reduced-order models with an effecient parallelization based on MPI. The implementation is tested on models of sizable state spaces using a high-performance computing node on Amazon Web Services, showing favorable scalability.

## 1. Introduction

The basic building blocks of living things are single cells, in which important biochemical processes occur and are regulated. There is increasing evidence that these chemical processes can lead to diverse outcomes, even in a population of cells with the same genetic makeup [19]. An important factor contributing to this variability is intrinsic noise due to random occurences of biochemical reactions. Such noise is usually modelled using a class of continuous time Markov chain models called stochastic reaction networks (SRNs) which treat molecular copy numbers of chemical species as random integers. These SRNs have been applied in various quantitative studies of real biological systems [45, 43], and can potentially be used for guiding optimal design of new experiments [20].

Many popular methods for the numerical study of SRNs are based on drawing sample paths from the underlying Markov process. The earliest example of the trajectorial approach is Gillespie’s algorithm [26], which remains the most popular simulation method for SRNs. There are also works on more effecient variants of Gille-spie’s algorithm [25], as well as approximate sampling approaches such as tau-leaping algorithms [27, 47, 10, 9, 3]. A major drawback of such Monte Carlo simulations is that convergence rate scales only as *O*(*n*^−1/2^), where *n* is the number of samples. As a consequence, the ease and low cost of simulating single paths may be offset by the need to draw a large number of samples to compute reliable estimates for quantities of interest such as moments and event probabilities. Furthermore, the accuracy of these estimates can only be assessed in terms of confidence intervals. There are goal-oriented approaches that reduce the variance and cost of Monte Carlo estimates such as multilevel sampling approaches [4, 36]. On the other hand, various numerical techniques aim instead at computing summary statistics of the probability distribution function, such as the mean and variance of molecular copy numbers at specific times. When the reaction networks consist of only monomolecular reactions with mass action kinetics, the moments of the copy numbers could be captured exactly by a finite system of ordinary differential equations (ODEs). In general, however, the lower-order moments depend on higher-order moments and an infinite number of moment equations is required to capture just the mean dynamics. There has been much work on moment closure techniques to approximate these inifinite systems by a finite sets of equations corresponding to low-order moments [33, 2, 35, 39, 49, 48]. While these methods can work effeciently and accurately for some SRN models, the error induced by the truncation technique is generally unknown. To circumvent this issue, there are recent works on computing bounds for the transient and stationary moments using semidefinite programming instead of closure [34, 17, 16].

This paper concerns a different approach that seeks to compute directly the time-dependent probability distribution of the process by solving the chemical master equation (CME). The finite state projection (FSP) [42] is a well-known representative of these approaches. In a nutshell, this method consists of truncating the infinite state space of the underlying Markov chain into a finite subset of states, effectively reducing the infinite-dimensional system of ODEs into a finite problem that is amenable to numerical integrators. In contrast to simulation and moment approaches, the FSP provides guarantee of accuracy with a deterministic error bound that can be easily computed from the solution of the truncated system [42]. The major drawback of the FSP is that the cost of solving the truncated system could grow prohibitive when there are many species or when the probability landscape is spread out over large number of states. Much has been done on improving algorithimc aspects of the FSP such as the selection of the state space [41, 7, 56, 8] and alternative tensor formulations [32, 53]. However, most of these algorithms were only implemented in serial, which limited the complexity of the models that could be solved with the FSP approach. There have only been very few works that discuss how to scale up the FSP on modern high performance computing platforms such as multiple-core cluster nodes [58, 52] or graphic processing units [38]. These early studies did not address the problem of exploring and updating the large state space of the CME in parallel, and the resulting implementations were only tested on truncated problems of at most a few million states.

In this paper, we introduce a novel adaptive parallel implementation of the FSP with particular focus on parallel exploration of the state space and the dynamic load-balance of the computations. We extend and parallelize a variant of the FSP described in [44], which makes use of an economical projection of the full CME state space that only expands when needed, using the proven error bound in [42] as the adaptation threshold. Most importantly, we parallelize the time-consuming task of managing the states of the FSP by distributing the storage, exploration, and migration of the states over all processors. This is a major contrast to earlier MPI-based implementations of the FSP, which either computed the state space offline in serial [58], or communicating the states in a all-to-one fashion [52]. We parallelize the numerical linear algebra computations involved in the FSP algorithm based on objects and routines from the library PETSc [5, 6], while intefacing with load-balancing approaches provided by Zoltan and ParMetis [14, 31]. Our solver can tackle CMEs with both time-invariant and time-varying propensity functions. In particular, we provide for the first time a parallel implementation of the Krylov-based Incomplete Orthogonalization Procedure [55, 24] for solving models with time-invariant propensities. We also provide interfaces into the high performance ODEs solution library SUNDIALS [30] for the solution of the general CMEs with time-varying propensities.

The paper is organized as follows. In section 2 we review the stochastic reaction networks modeling approach that gives rise to the CME, and the basic principles of the FSP algorithm. In section 3 we describe a novel adaptive FSP method and review approaches for solving the resulting reduced systems. In section 4, we discuss the parallel implementation of this adaptive FSP. Numerical experiments are discussed in section 5, where with 36 cores of an Amazon Web Services computing node we solve in a few minutes a demanding problem with 23 million states that would have taken hours to solve with a single CPU. Section 6 summarizes the paper’s findings and discusses future work.

The default notational convention is as follows. For a vector ***v***, we denote its *i*-th entry by either the subscripted notation [***v***]_*i*_ or the MATLAB-like notation ***v***(*i*). Similarly, an entry of a matrix ***A*** could be denoted as either [***A***]_*i,j*_ or ***A***(*i, j*). We denote by nnz (***v***) and nnz (***A***) the numbers of nonzero entries in respectively ***v*** and ***A***. We use the boldface notation ***p*** to denote the probability distribution obtained from solving the CME. The notation *n*_proc_ is reserved for the number of processors. For a set *S*, we denote by |*S*| the number of elements in *S*.

## 2. Background

### 2.1. Stochastic reaction network models of gene expression

Consider a chemical reaction network with *N* molecular species with *M* reaction channels. The state of the chemical network is the vector ***x*** ∈ ℕ^*N*^ of copy numbers of different molecular species. Each reaction event, such as transcription of a new mRNA molecule, incurs a change in the state that is captured by a stoichiometric vector ***ν***_*k*_ ∈ ℕ^*N*^. Given the current state ***x***, the process can transit only to states of the form ***x*** + ***ν***_*k*_, *k* = 1, …, *M*. The time-dependent state vector is modeled as a continuous-time Markov chain {***X***(*t*)}_*t*≥0_ with transition rates given by the propensity functions *a*_*k*_(*t*, ***x***). From a current state ***x***, the chain can make jumps to *M* neighboring states ***x*** + ***ν***_*k*_ with the infinitesimal probability *a*_*k*_(*t*, ***x***).

The chemical master equation (CME) describes the dynamic of the probability distribution of the state ***X***(*t*). In particular, given the initial state ***X***(0) with known distribution, let *p*(*t*, ***x***) be the probability of the event [***X***(*t*) = ***x*** | ***X***_0_], then the CME has the form

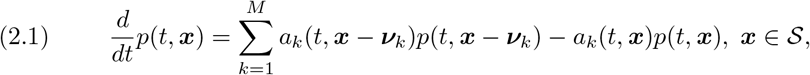

where 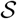 is the set of all reachable states.

We restrict our attention to models where the propensity functions are separable in the form

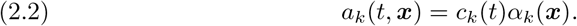

This assumption is valid for any systems with time-invariant propensities, and for systems with time-varying propensities that follow mass-action kinetics.

Since the state space 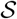 is discrete, we can treat the probability distribution of ***X***(*t*) as a probability vector ***p***(*t*). Define the transition rate matrix ***A***(*t*) with entries enumerated by the states as

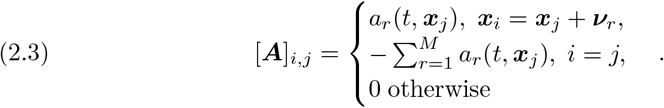

Equation (2.1) could then be equivalently formulated as an initial value problem that takes the form

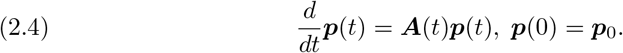

Next, we discuss how the CME could be solved numerically using the finite state projection.

### 2.2. The Finite State Projection algorithm

The finite state projection (FSP) [42] is a systematic approach to approximate the solution of the CME using a vector with finite support. In its simplest formulation, the FSP consists of the truncation of the infinite state space 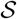 into a finite set 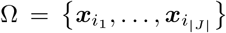. Let *J* = {*i*_1_, …, *i*_|*J*|_} be the corresponding set of indexes. For simplicity and without loss of generality, let us assume that *J* = {1, 2, …, |*J*|}.

The FSP first solves a finite system of the form

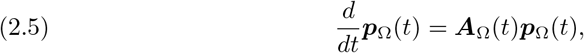

where

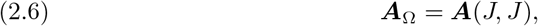

and

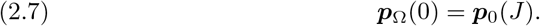

The solution of the finite system (2.5) provides the approximation to the true CME solution on Ω, that is,

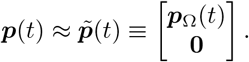

The FSP could be viewed as an aggregation method that turns all transitions to outside of Ω into an absorbing state (Fig. 1). The probability of this sink state provides the exact approximation error [42, Theorem 2.2] in terms of the one-norm,

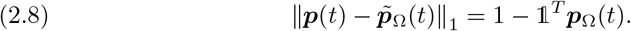

**FIG. 1.**
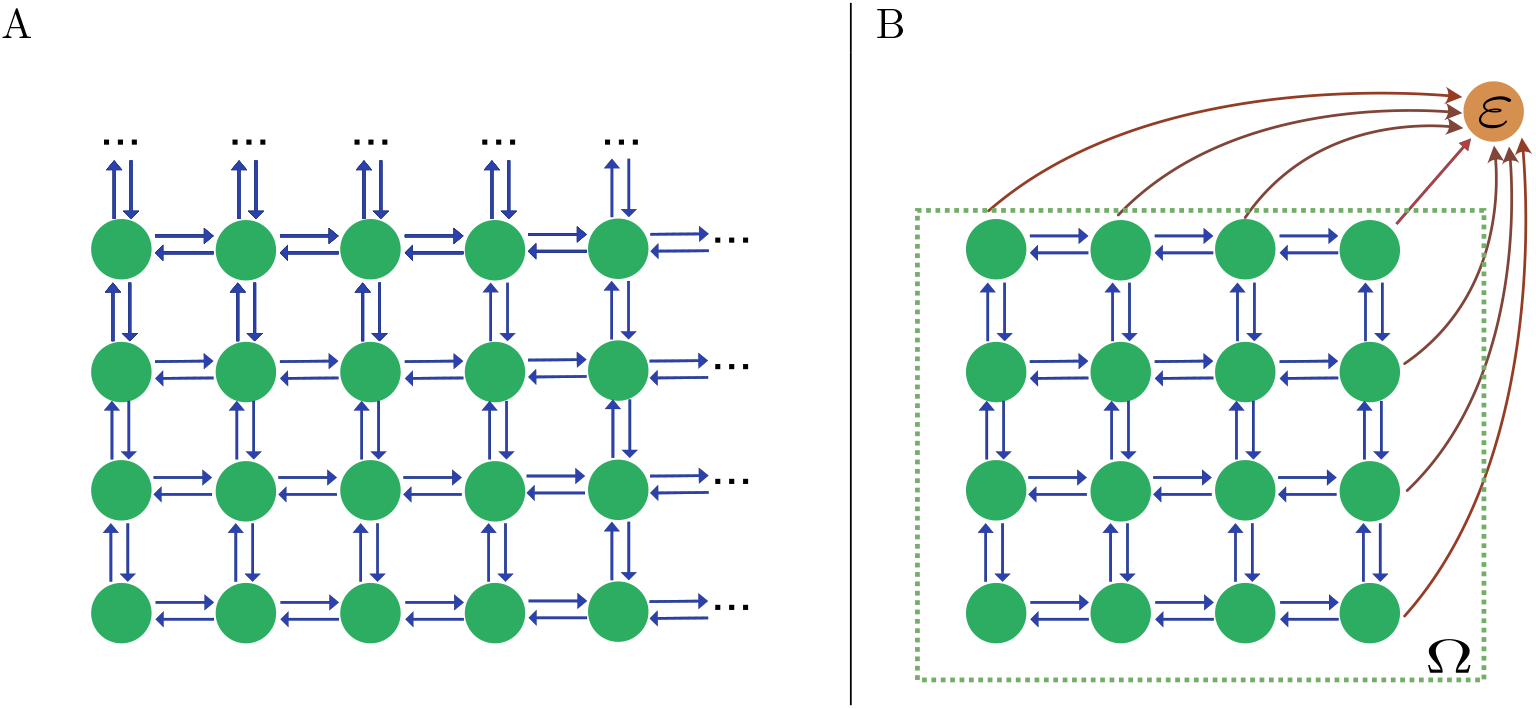
Schematic of the Finite State Projection algorithm. (A): The original state space of the CME, which could have infinitely many states. (B): The truncated state space, where the states outside a finite subset (Ω) are lumped into a single absorbing state (ε). The probability of ε is found simply by subtracting to one the sum of probabilities of the remaining states (eq.(2.8)).

For all models encountered in practice, the error bound *g*(*t*) decreases monotonically as Ω expands [42]. Under some regularity conditions on the propensity functions, it could be shown that 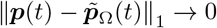 as 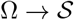 [22].

## 3. Numerical schemes

### 3.1. Adaptive Finite State Projection with multiple sink states

We use finite state sets Ω that consist of reachable states that satisfy a set of constraints [44]

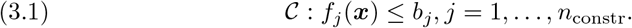

where *b*_*j*_ are the upper bounds that are changed in an adaptive way as the FSP integration progresses. Let 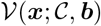 denote the set of constraints determined by 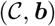 that is violated by a state ***x*** ∈ Ω, and let 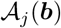 denote the set of states that satisfy the *j*-th constraint. Furthermore, let 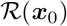 denote the set of states that can be reached from the initial state ***x***_0_. The finite state set for the FSP is thus

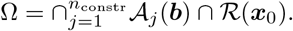

We then approximate the Markov process of the CME with a finite Markov process with the augmented state space 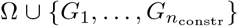, where each *G*_*ℓ*_ (*ℓ* = 1, …, *n*_constr_) is a sink state that absorbs transitions from Ω into states that violate the *ℓ*-th constraint. Fig. 2 presents an example of the truncated state space based on a set of three constraints.

**FIG. 2.**
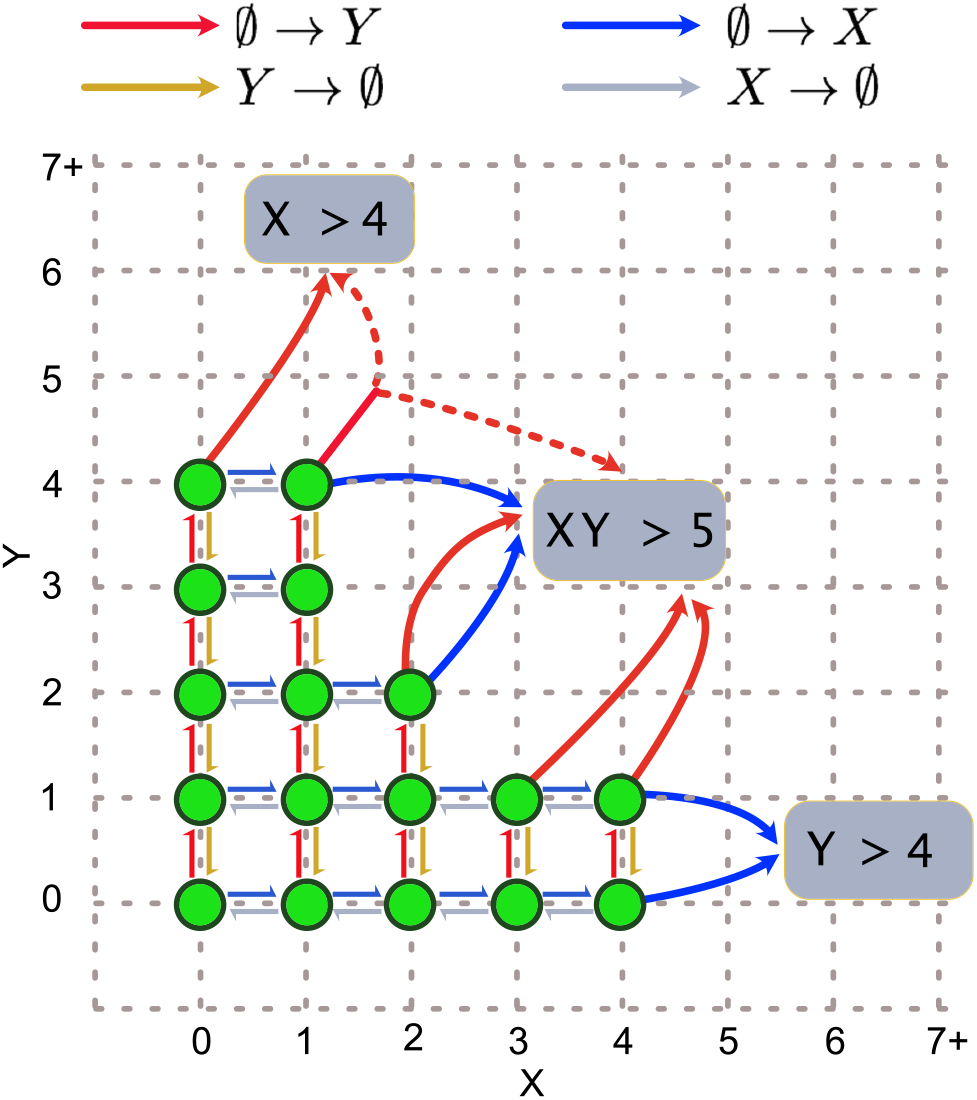
Example of the finite state projection with multiple sink states. We consider a simple two-dimensional model with four reactions ∅ → X, X → ∅, ∅ → Y, Y → ∅. The sink states correspond to three constraints: (ε_0_): [*X*] ≤ 4, (ε_1_): [*Y*] ≤ 4, (ε_2_): [*X*][*Y*] ≤ 5. The arrows represent transitions from each state. Transitions leading to states that violate the j-th constraint will be directed to ε_j_. Note that a reaction may bring one state into violating different constraints simultaneously (state (1, 4) in the graphics), in which case the transition rate is divided equally among the receiving absorbing states.

Assuming the augmented state space is enumerated so that the sink state *G*_*ℓ*_ has index |Ω| + *ℓ*, the time-varying transition rate matrix 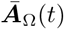 of this Markov process could be decomposed into

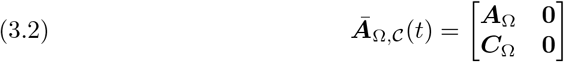

where ***A***_Ω_ captures the transitions between states in Ω and 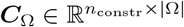 captures the transitions into the absorbing states. Here, if a transition ***x*** → ***x***+***ν***_*r*_ violates more than one constraints then we divide the rate *a*_*r*_(*t*, ***x***) equally into the corresponding absorbing states. The nonzero entries of ***A***_Ω_ and ***C***_Ω_ are given by

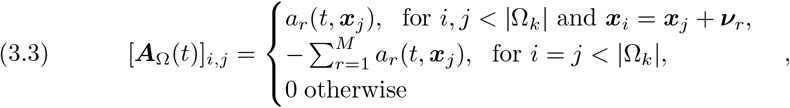

and

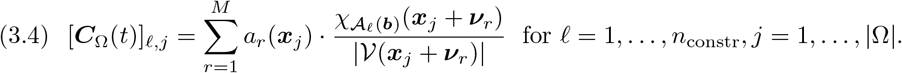

The FSP approximation becomes

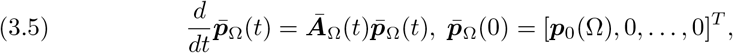

where the probability vector 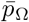 is partitioned into

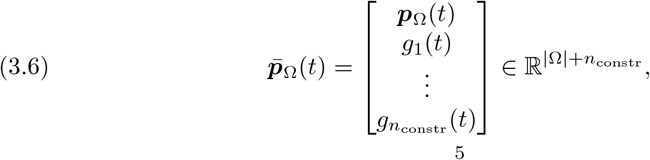

with the last *n*_constr_ entries of 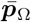 tracking the probabilities of the absorbing states.

The advantage of the multi-sink formulaion (3.6) to the original one-sink FSP is that we know which sink state attracts the most probability and we can selectively relax only their corresponding constraints for the next iteration. As a consequent, the finite state set Ω only needs to expand on the directions where the probability mass leaks the most, allowing for a more flexible and economical truncation.

Taking this one step further, we divide the whole time interval [0, *t*_*f*_] into subintervals with timesteps 0 ≔ *t*_0_ < *t*_1_ < … < *t*_*K*_ ≔ *t*_*f*_, and devise a multiple-time-interval formulation [7, 56]. At each subinterval [*t*_*k*_, *t*_*k*+1_) we solve a finite system of the form

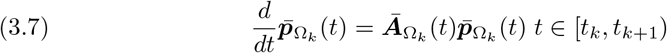

Each solution vector 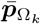 has the partitioning (3.6), with the absorbing state probabilities *g*_*j*_(*t*), *j* = 1, …, *n*_constr_ tracking the accumulated probability mass that are loss by various FSP approximations. Let 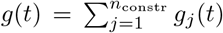, then *g*(*t*) is an upper bound on the accumulated approximation error. We require that the maximum error acummulated at the absorbing state satisfies

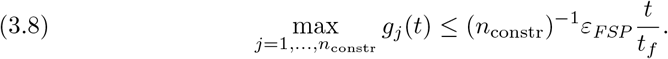

Given the current time step *t*_*k*_, we let the ODE integrator advance the solution until either reaching final time *t*_*f*_ or an itermediate timepoint *t*_*k*+1_ where the inequality(3.8) starts to change direction.

If the integration halts at an intermediate timepoint due to the error control (3.8), we enlarge the FSP constraint bounds *b*_*j*_ ← (1 + *δ*)*b*_*j*_, *j* = 1, …, *n*_constr_, giving rise to a new set of relaxed constraints 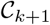. We then expand from Ω_*k*_ into states that satisfy these more relaxed constraints, giving rise to the expanded set Ω_*k*+1_ that constitutes the FSP approximation of the next time interval. The recommended scaling factor in our code is *δ* ≔ 0.2, though this could be changed by the user.

The pseudocode in Algorithm 3.1 describes the prototype of the adaptive Finite State Projection algorithm we just discussed. From there, we see that an implementation of the algorithm breaks down to three major groups of tasks: the dynamic management of the finite state subset Ω_*k*_, the generation of the time-dependent matrix 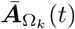 and the matrix-vector multplications at each time step, and the advancement of the resulting finite system of ODEs.

**Algorithm 3.1.**
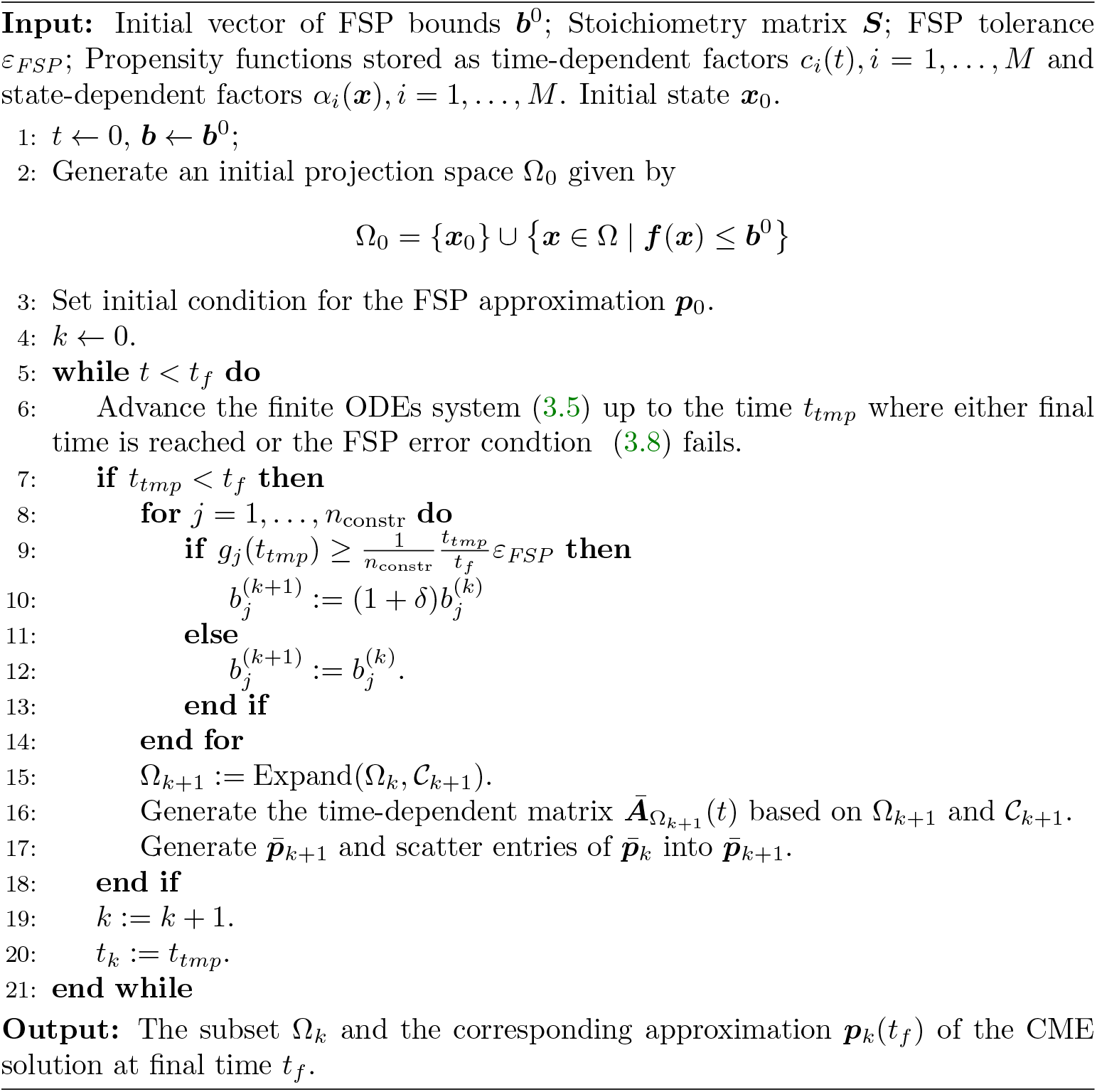
Multiple-sink-state Finite State Projection

### 3.2. Adaptive Krylov integrator with event detection for the time-invariant propensities

When all propensities are time-invariant, solving the finite systems (3.7) amounts to computing the action of the matrix exponential,

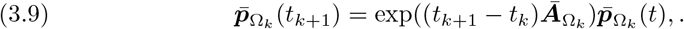

A popular approach for evaluating these expressions that are particularly well-suited for large-sparse matrices arising from the FSP are Krylov subspace techniques [50], which have been incorporated in many serial CME solvers [7, 51, 8].

To simplify notations, let 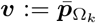 and 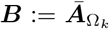. The Krylov-based approach starts by generating a pair of matrices 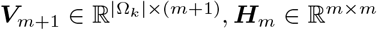, such that the columns of ***V***_*m*+1_ ≔ [***v***_1_ ***v***_2_ … ***v***_*m*+1_] form a basis for the Krylov subspace

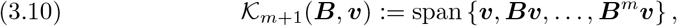

and that the following relation is satisfied,

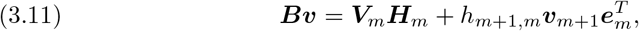

where ***V***_*m*_ ≔ ***V*** (:, 1 : *m*), *h*_*m*+1,*m*_ ≔ ||***Bv***_*m*_||_2_, and ***e***_*m*_ = [0 … 01]^*T*^ ∈ ***R***^*m*^. Supposing that these conditions are met, the pair (***V***_*m*_, ***H***_*m*_) can then be used to form an approximation for the matrix exponential operator as

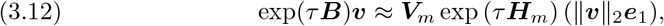

where ***e***_1_ = [1 0 … 0]^*T*^ ∈ ℝ^*m*^. The exponential of the small dense matrix *τ* ***H***_*m*_ in the above expression is usually computed using the diagonal Padé method [50].

There are many ways to generate the matrices ***V***_*m*+1_ and ***H***_*m*_ that satisfy the two conditions described above. The well-known implementation in the package Expokit [50], for example, uses the full orthogonalization method (FOM), which ensures that ***V***_*m*+1_ is an orthogonal matrix, making the approximation (3.12) equivalent to an orthogonal projection onto the corresponding Krylov subspace. The FOM, however, requires modified Gram-Schmidt sweeps whose cost grows quadratically with the Krylov dimension *m*. This becomes prohibitive for large input vectors ***v***. There are recent works that propose the alternative use of Incomplete Orthogonalization Procedure (IOP) [55, 24] that trade the cost of full orthogonalization for a slight increase in matrix-vector multiplications and larger Krylov dimension. This is the approach that we implement in this paper.

In contrast to the FOM, the IOP requires only that each vector ***v***_*j*_ be orthogonal to the preceding *q* vectors ***v***_*j*−*q*_, …, ***v***_*j*−1_, where *q* ≥ 1 is a user-prescribed orthogonalization length (the case *q* = *m* reduces to the classic Arnoldi procedure). When implemented using an adaptive strategy based on the *phipm* algorithm of Niesen and Wright [46], the IOP with *q* ≔ 2 outperform the traditional Arnoldi procedure on several numerical benchmark problems drawn from computer and biochemical systems [55] and shallow water equations [23]. The details of this strategy could be found in [55]. We discuss here only the aspects that are relevant to the parallel implementation of the IOP, and also note that a parallel implementation for the FOM with variable Krylov subspace dimension has been previously proposed [37].

Similar to the serial implementations [23, 55], we break down the matrix exponential (3.9) into a step-by-step integration scheme that advances through timesteps 0 ≔ *t*_1_ < *t*_2_ < *t*_*J*_ ≔ *T*. The intermediate solutions are given by the Krylov approximations

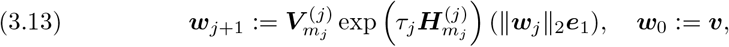

where, similar to (3.12), 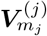 and 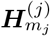 are generated from the IOP applied to the matrix ***B*** and the vector ***w***_*j*_. Note that the stepsize *τ*_*j*_ ≔ *t*_*j*+1_ − *t*_*j*_ and the Krylov subspace dimension *m*_*k*_ are not fixed, but determined adaptively during the integration. After a successful step at time *t*_*j*_, we either keep the Krylov dimension fixed while using a new stepsize *τ*^*new*^ for the next step, or keep the current stepsize but use a new value *m*^*new*^ for the Krylov basis size. These stepsize and basis size values are determined from a criteria that ensures that a posteriori error estimate of the local truncation error remains below a tolerance (see [55] for details of error estimation and control). The decision to vary which quantity (*τ*_*j*_ or *m*_*j*_) is based on a guess of the future integration cost associated with each choice.

We estimate the local cost per CPU for advancing with a stepsize *τ* and basis size *m* is given by [46, 23, 55]

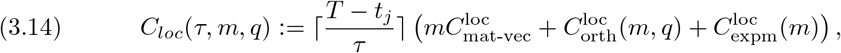

where 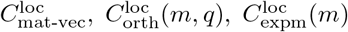 are the on-processor costs, measured in terms of the number of FLOPs, for matrix-vector multplications with ***B***, orthogonalization cost for basis size *m* and IOP parameter *q*, and the evaluation of the small matrix exponential of size *m* × *m* with Pade technique.

These costs are estimated in the same way as in [53], but with the global sizes of the matrix and vectors involved replaced by their local dimensions. Once these local costs are computed for each pair (*τ*_*j*_, *m*^*new*^) and (*τ*^*new*^, *m*_*j*_), we make a call to `MPI_Reduce` to get the maximum cost *C*_*glob*_(*τ, m, q*) over all processors. We then choose to change dimension if *C*_*glob*_(*τ*^*new*^, *m, q*) > *C*_*glob*_(*τ*, *m*^*new*^, *q*) and to change the stepsize otherwise. We choose to use the maximum over local costs, rather than the sum as in a previous parallel adaptive Krylov implemention [37], since parallel computational time depends more on the processor with the heaviest workload than on the total workloads over all processors.

## 4. Parallel implementation

### 4.1. Parallel management and dymanic expansion of the state space

The truncated state set Ω is partitioned into Ω^(*j*)^ ⊂ Ω, each separately owned by a processor. A state 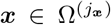 held by processor *j*_***x***_ has a local index *i*_loc_(***x***) ∈ {1, …, |Ω^(*j*)^|}. To manage and retrieve information about the states in a scalable way, we employ the Distributed Directory (DD) in the Zoltan library [14]. The DD object is essentially an MPI-based hash table that take multi-dimensional vectors of integers as keys. The data entry for each key 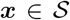 consists of a pair (*j*_***x***_, *i*_loc_(***x***)) that stores the rank of the owning processor and the local index of ***x***. Each call to the lookup and insertion routines must be done collectively by all processors.

Let Ω_*k*_ be the state set at timestep *k* of the Finite State Projection. In order to expand Ω_*k*_ into a new set Ω_*k*+1_ determined by the constraints 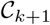, we use an iterative breadth-first search algorithm. Let *∂*Ω denote the set of states on the “boundary” of 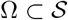, that is,

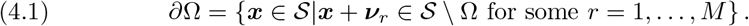

The serial state space expansion scheme starts at the current state set Ω_*now*_ ≔ Ω_*k*_. We then compute the set 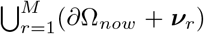 and thin it down to a subset *B* that consists only states satisfying 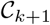. We then annex *B* to Ω_*now*_, and the process repeats until all reachable states that satisfy 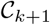 have been explored.

Our parallelization of this scheme is described in algorithm 4.1. On line 3 of the pseudocode, we partition *∂*Ω_*now*_ into roughly equal parts that are distributed to individual processors. This helps improve the load-balance of the exploration step on line 5-7. The distribution step employs a fast and simple partitioning scheme (called `BLOCK` in Zoltan) that distributes roughly equal numbers of states to the processors. Then, the exploration of new states on line 5-7 is done concurrently on all processors without the need of inter-processor communication. We then annex the states discovered thus far to the global state set, while making sure to not add redundant states by checking with the Distributed Directory.

**Algorithm 4.1.**
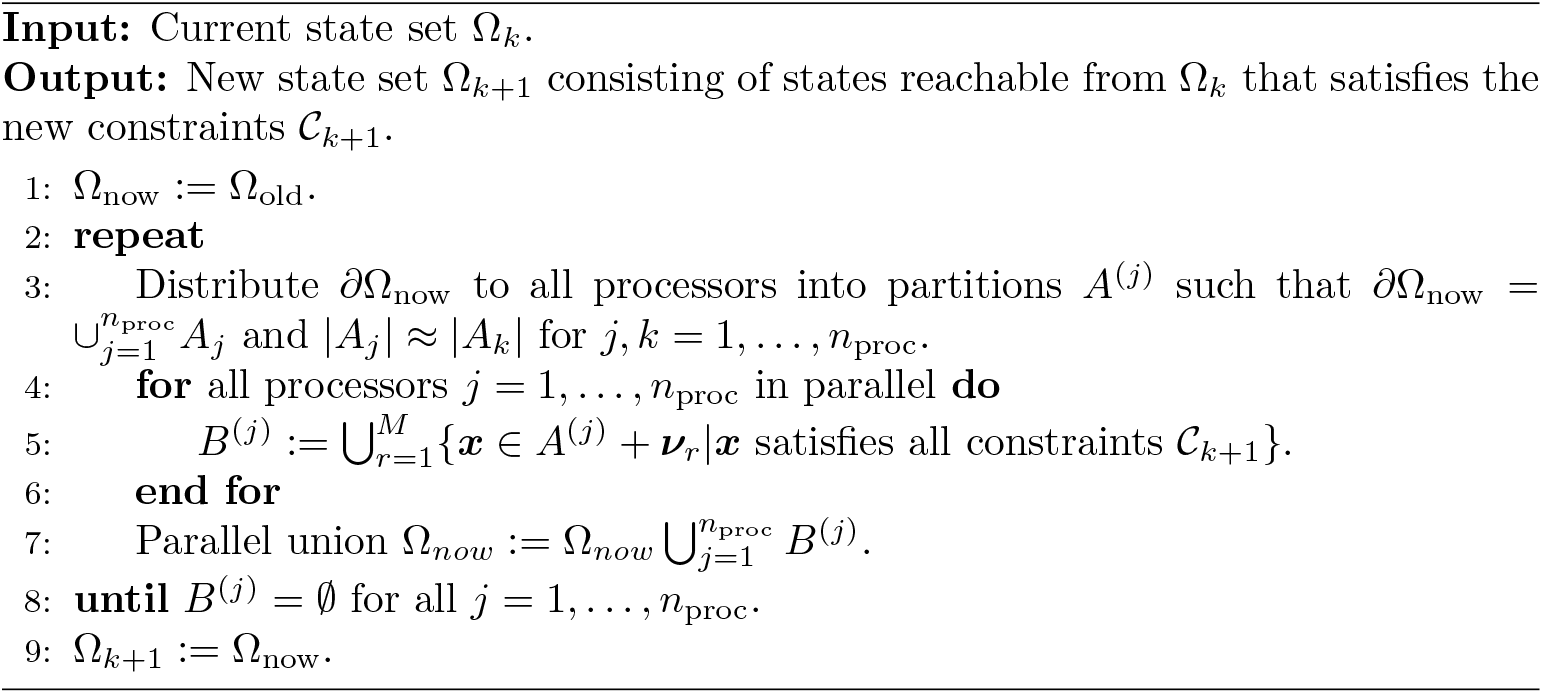
Parallel expansion of the finite state subset.

### 4.2. Parallel matrix object

Let 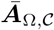 be the time-dependent transition rate matrix associated with a state set Ω and the constraint set 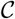. Recall that we can partition 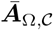 into ***A***_Ω_ and 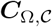, defined respectively in eq. (3.3) and eq. (3.4). Recall separability assumption (2.2) on the propensities, and let 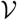 denote the set of reactions *r* for which *c*_*r*_(*t*) varies with the time variable *t*. These matrices have the decompositions

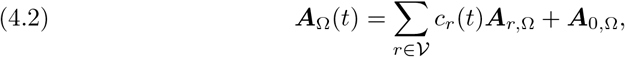

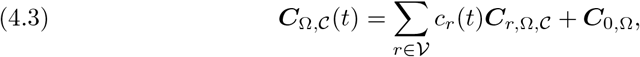

where 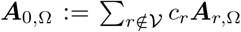 and 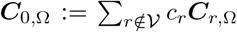 captures the time-invariant part of the matrices ***A***_Ω_ and 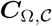.

As illustrated in fig. 3, each ***A***_*r*_ is distributed row-wise into the processors as done in PETSc’s parallel Mat object [5], while the (very thin) ***C***_*r*_ matrices are distributed column-wise. The column-wise partitioning of ***C***_*r*_ means that no off-processor elements of the input vector is needed for the parallel matrix-vector multiplication with 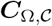, while the update of the global output vector requires only *O*(*n*_constr_) messages. PETSc offers a handful of different storage schemes for the on-processor portions of the sparse matrix. We found in our tests that storing the FSP matrix with the `MATMPISELL` format [57] yields better performance than the default compressed sparse row (CSR) format. This is also the matrix format that will be used in the numerical tests below.

**FIG. 3.**
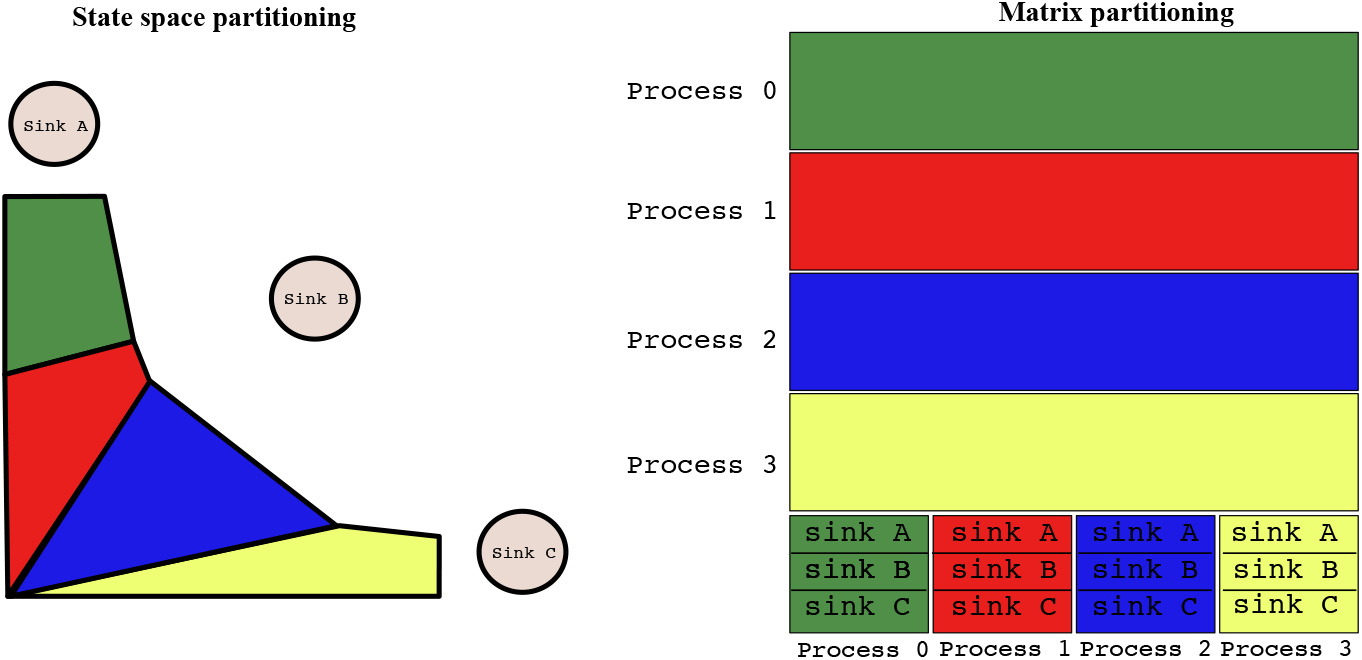
Relationship between state partitioning and the partitioning of the FSP-reduced transit rate matrix 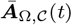 (eq. (3.5) in the main text). The submatrix corresponding to transitions between states is distributed row-wise to the processes, while the submatrix corresponding to transitions into sink states is distributed column-wise.

### 4.3. Dynamic load-balancing approaches

Since the adaptive FSP adds states to the projection space after every time step, it is important to redistribute the states and their associated data after expansion, so that the subsequent computational work loads are evenly distributed among the processors, while keeping inter-processor communication minimal. In addition, the redistriution scheme must also take into account the cost of data migration. While the true computation and communication cost is generally hard to quantify, several approximations and heuristics based on graphs and hypergraphs have been shown to yield good performance in practice [11, 31, 12]. We specifically interface our state space routines with the Graph-based adaptive repartitioning algorithm of ParMETIS [31] and the hypergraph-based PHG algorithm of Zoltan [14, 12], and also to a simple strategy that only seeks to distribute CME states equally among processors.

### 4.4. Code availability and Python interface

All the numerical techniques and algorithms discussed have been implemented as a C++ library ^1^ that can also be used in Python using a Cython-based wrapper ^2^. The features of the C++ programming language allow us to write our codes in an object-oriented way, with an eye on reusability and extensibility. Specifically, the basic building blocks of the presently discussed FSP variant (as well as future developments) are the three object classes `StateSetBase, FspMatrixBase`, and `OdeSolverBase`. These objects correspond to the basic data structures and methods that are common in all FSP methods. In particular:

1. The `StateSetBase` class manages the subset of CME states. It provides basic methods for state lookup and insertion, as well as parallel state partitioning and redistribution (via interface to Zoltan [14]). It has a virtual method `Expand` to be specified by more specialized classes to implement the specific state space expansion technique. In particular, we have a child class `StateSetConstrained` that implement the nonlinear constraints-based state expansion technique mentioned in section 4.1. Other methods for expanding the state space [51, 56] can also be similarly developed by specifying only the `Expand` method.
2. The `FspMatrixBase` class manages the time-dependent truncated transition rate matrix of the CME. This class provides the basic method to generate the state transition rate matrix resulted from FSP truncation as defined in eq. (3.3) (but not the multi-sink components (3.4), which we leave to a child class named `FspMatrixConstrained`, which works with `StateSetConstrained` and which generates the additional absorbing state components needed for the multi-sink FSP solver).
3. The `OdeSolverBase` class provides basic methods to solve the truncated linear ODE systems (3.7). The main class data is the pointer to an instance of `FspMatrixBase`, and a pointer to a function that perform the check for the FSP criteria (2.8). This means that the ODE solution can halt midway if it finds the FSP error to exceed the prescribed tolerance, and return to the FSP solver to request for an expanded state space. The specific ODE solver used is provided in child classes that provide interfaces to PETSc’s TS module [1] and SUNDIALS [30], as well as implementing the Krylov-based IOP technique discussed in section 4.

The multi-sink FSP approach we introduce in this paper is implemented in a class called `FspSolverMultiSinks`, which is based on the composition of the objects discussed above. We note that parallel implementations of other FSP-based algorithms [7, 51, 56, 8, 29] can be similarly benefited from reusing the base classes we mentioned above.

## 5. Results

In this section, we present numerical tests of our parallel FSP implementation on three different stochastic chemical kinetics models. The aim of these tests is to demonstrate how different algorithmic choices affect the runtime and scalability of the solver on specific problems. All tests are run on a c5.18x instance of Amazon Web Serivces. This node consists of two sockets, each of which houses 18 Intel Xeon Platinum 8124M CPU cores that operate at 3.00GHz per core. We use PETSc 3.11.3 and Zoltan library that were compiled against OpenMPI 4.0 with the GNU C/C++ compilers. In addition to default build settings, the PETSc library is compiled with the option to enable AVX-512 kernels [57], which Intel Xeon processors support. The execution of the parallel programs are done via a call to `mpirun` with the options `--bind-to-core` and `--map-by-socket` which binds each process to a physical core and which spreads the processes as evenly as possible across the NUMA nodes. Our codes are written in an object-oriented style in C++ and are compiled using OpenMPI compiler wrapper of GNU.

### 5.1. Repressilator

We first consider a three-species model inspired by the well-known repressilator gene circuit [18]. This model consists of three species, TetR, *λ*cI and LacI, which constitute a negative feedback network (Table 1). We solve the problem up to time *t* = 10 (abitrary time unit) using the FSP tolerance of 10^−4^, starting from a point mass measure concentrated at ***x***_0_ = (TetR, *λ*cI, LacI) = (20, 0, 0). We let the state space grow adaptively and use the Adaptive Repartitioning algorithm of ParMetis [31] to dynamically re-balance the state space across processors.

**TABLE 1.** Reactions and propensities in the repressilator model. ([X] is the number of copies of the species X.)

|  | reaction | propensity | parameters (arbitrary unit) |
| --- | --- | --- | --- |
| 1. | $\emptyset \rightarrow \text{TetR}$ | $k/(1 + a[\text{LacI}]^b)$ | $k = 100, a = 20, b = 6$ |
| 2. | $\text{TetR} \rightarrow \emptyset$ | $\gamma[\text{TetR}]$ | $\gamma = 1$ |
| 3. | $\emptyset \rightarrow \lambda\text{cI}$ | $k/(1 + a[\text{TetR}]^b)$ | |
| 4. | $\lambda\text{cI} \rightarrow \emptyset$ | $\gamma[\lambda\text{cI}]$ | |
| 5. | $\emptyset \rightarrow \text{LacI}$ | $k/(1 + a[\lambda\text{cI}]^b)$ | |
| 6. | $\text{LacI} \rightarrow \emptyset$ | $\gamma[\text{LacI}]$ | |

We compare the performance of four algorithmic variants based on the choice of the FSP shape and on whether the FSP grows dynamically or is fixed at the beginning of the integration. For the choice of FSP shape, we can define the FSP state space is the customary way of choosing a hyper-rectangle defined by the constraints *x*_*i*_ ≤ *b*_i_, *i* = 1, 2, 3. However, the structure of the repressilator network suggests that we can exploit the negative correlation between species to truncate more states. In particular, we can add more nonlinear constraints of the form *x*_*i*_*x*_*i*+1_ ≤ *b*_*i,i*+1_, *x*_3_*x*_1_ ≤ *b*_3,1_(*i* = 1, 2) which stems from the intuition that more copies of a repressing species lead to fewer copies of the repressed species. This adds up to a total of six constraints as detailed in Table 2.

**TABLE 2.** Constraints on the FSP for the repressilator model.

|  | constraint function | initial bound |
| --- | --- | --- |
| 1. | $[\text{TetR}]$ | 22 |
| 2. | $[\lambda\text{cI}]$ | 2 |
| 3. | $[\text{LacI}]$ | 2 |
| 4. | $[\text{TetR}] \cdot [\lambda\text{cI}]$ | 44 |
| 5. | $[\lambda\text{cI}] \cdot [\text{LacI}]$ | 4 |
| 6. | $[\text{TetR}] \cdot [\text{LacI}]$ | 44 |

For the option the explore the FSP states, the adaptive variants choose a small set of states at the beginning of the integration and explore more states dynamically as explained in section 3. The static FSP variants solve the truncated FSP systems with a large set of states fixed for the whole integration time. The constraint bounds for the static variants is chosen as the final constraints ouput from the respective adaptive variants.

The marginal distributions computed from the FSP solution at the final time is given in Fig. 4, with no visible difference in computational outputs between different FSP options. On the performance side, within the shape choice of shape constraints, there is no difference between the adaptive and the static variants 6. However, since an appropriate state space size is generally difficult to know beforehand, it is more convenient to use adaptive FSP. For the FSP shape constraints, we see that using the extended set of constraints results in a smaller state space (Fig. 5A), which means that the truncated ODEs systems to be solved in these variants are smaller. The FSP variants that use these nonlinear constraints consistently reduce the average workload per processor (in terms of FLOPs) across the numbers of processors (Fig. 5B), and consequently the computational time (Fig. 6). In addition, we also record the ratio of the maximum vs minimum workload per processor (Fig. 5C). These ratios stay well below 1.1, indicating that the dynamic load-balancing methods were able to distribute the computation almost equally among processes. On the other hand, parallel effeciency degrades as the number of parallel processes increases, with all variants drop below 50 percent effeciency when executed on 32 processors. This is because the gain from dividing the problem across the fast Intel Xeon processors has been offset by the overhead of managing inter-process communication. Nevertheless, there is additional speedup in all major components of the code (Fig. 7) as we increase the number of processes. This is a relatively small problem that could be solved on one core in an order of minutes, so the gains from parallelization is expectedly moderate.

**FIG. 4.**
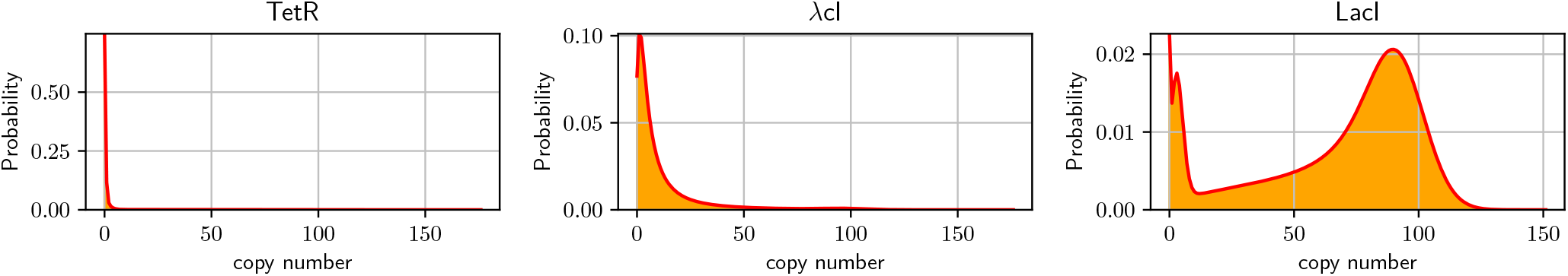
Repressilator example. Marginal distribution of TetR, λcI and LacI at time *t*_*f*_ = 10 minutes.

**FIG. 5.**
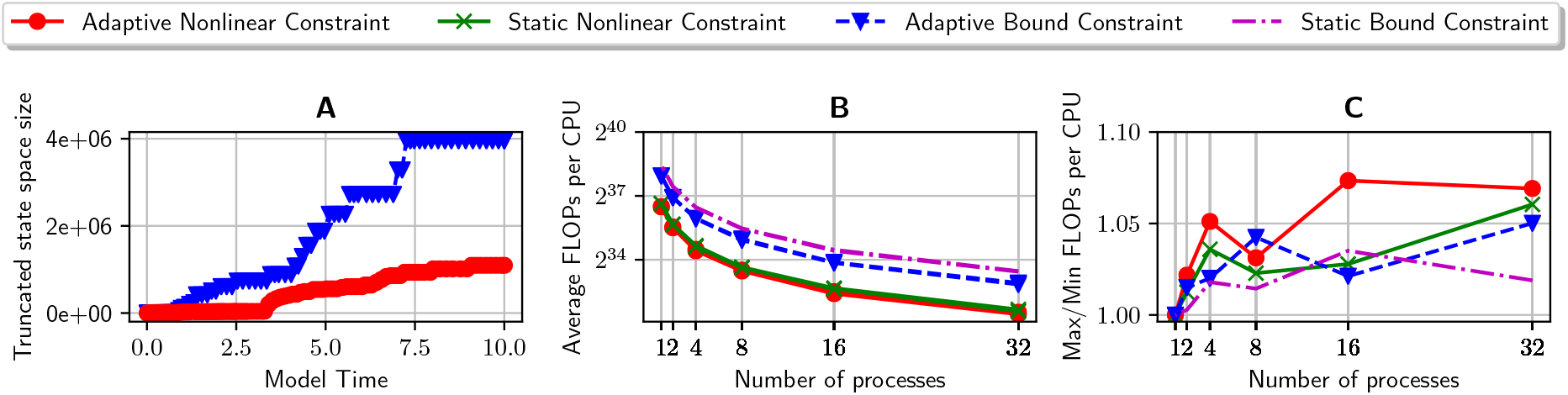
Workload in the numerical integration of the repressilator example. (A): Size of the FSP staet space over time for the adaptive variants, using the hyperectangular shape vs the nonlinear shape with additional constraints (cf. Table. 2). (B): average number of floating point operations (FLOPs) per process for four different choices of FSP shape and adaptivity. (C): The load-imbalance ratio, defined as the ratio between the maximum and minimum numbers of FLOPs per process, in the four algorithmic variants.

**FIG. 6.**
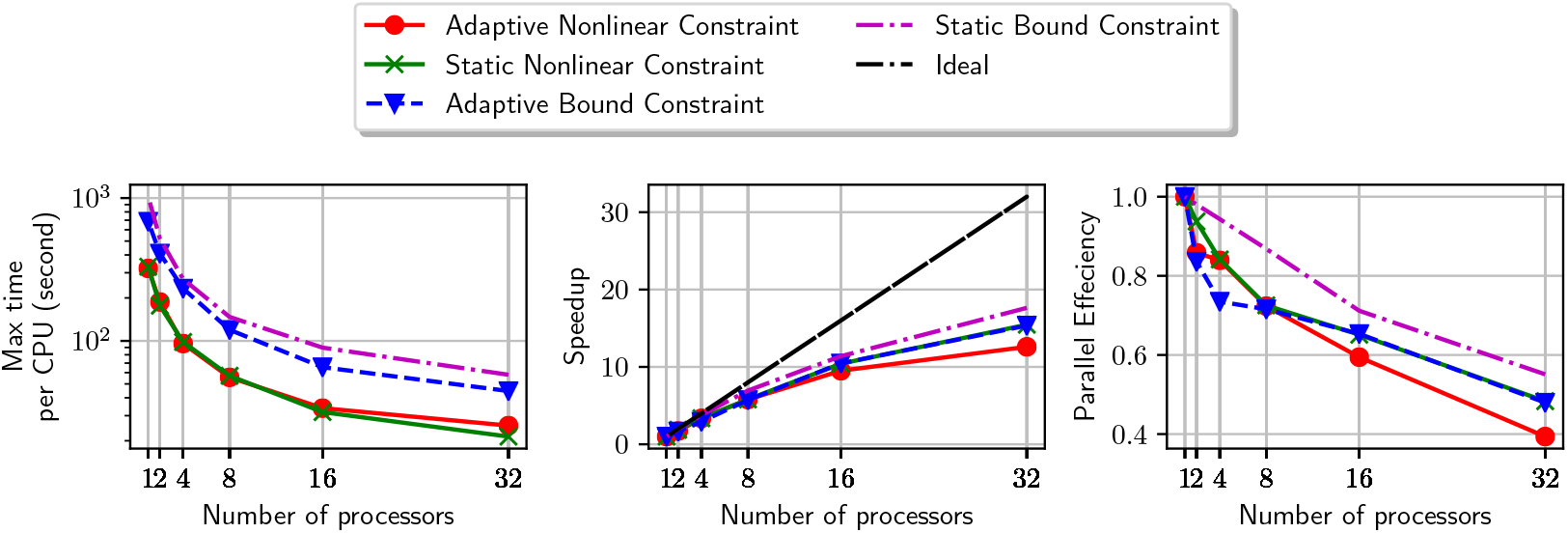
Computational time, speedup, and parallel effeciency in the parallel integration of the repressilator example. The four curves in each plot correspond to different choices for the FSP, using either the hyperectangular shape or the nonlinear shape with additional constraints (cf. Table. 2), with either adaptive or static FSP.

**FIG. 7.**
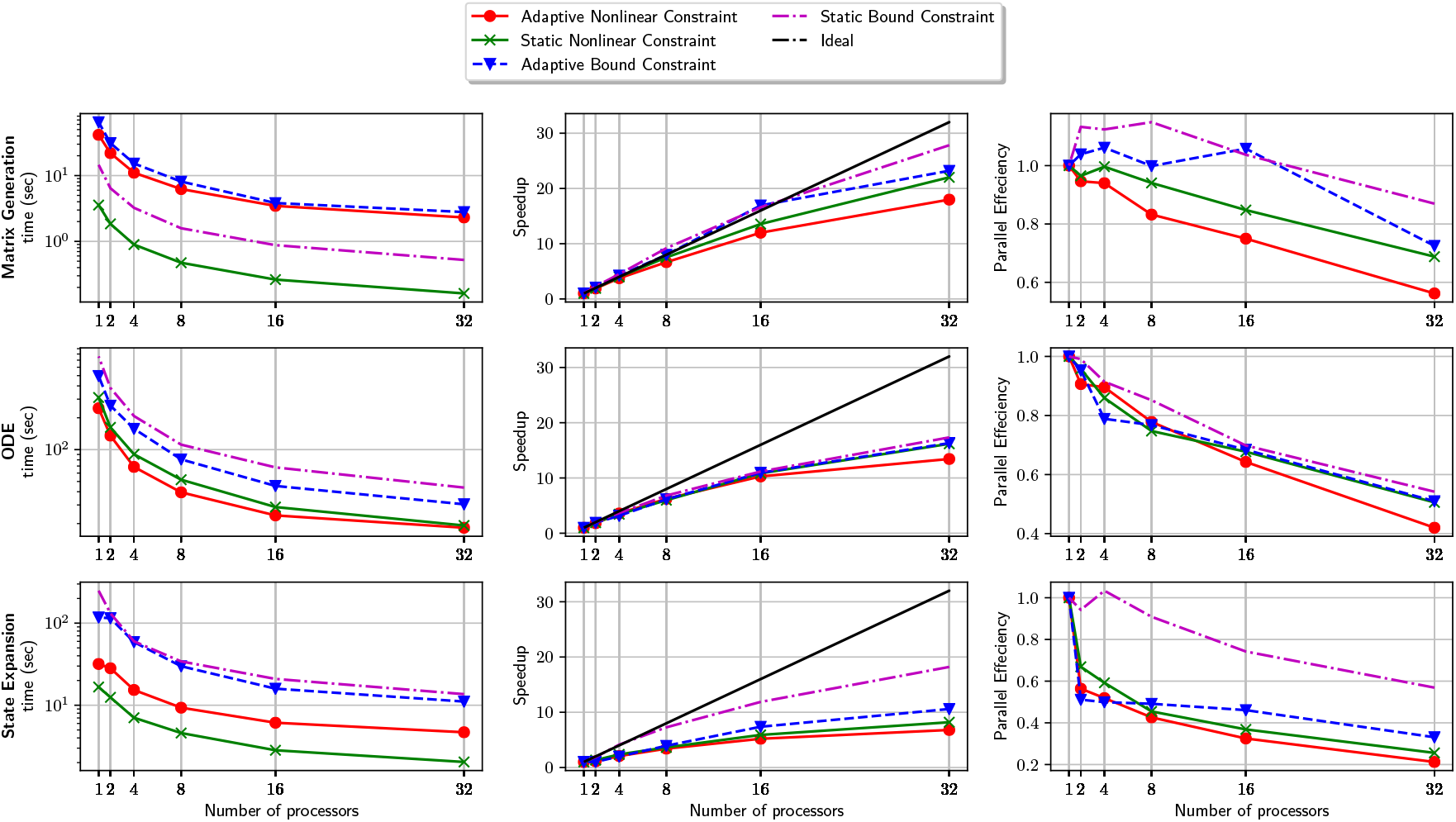
Time spent in the critical components in the parallel solution of repressilator example. These critical components include matrix generation (first row), solution of the truncated ODEs systems (second row), and state space expansion (third row). We compare the computational time, speedup, and parallel effeciency of four algorithmic variants, based on either using an adaptive of static FSP, and on using the nonlinear or hyper-rectangular constraints.

### 5.2. Six-species transcription regulation

We next consider a transcription regulation model introduced in [28]. In this model, the transcription-translation process of a protein M (monomer) is activated when a transcription factor D (dimer) binds to the DNA at a single site, but is repressed when D binds to both sites of the DNA’s promoter region. The propensities associated with second-order reactions are time-varying, due to the change in the cell’s volume (Table 3). This model results in a time-dependent distribution that spreads out over an increasing number of states over time, and is usually used as a benchmark problem for direct CME methods [7, 56, 51]. Most of the previous works only attempt at the time-invariant version of this model. We are only aware of the paper of Dinh and Sidje [15] that makes an attempt at directly solving the CME with time-varying propensities in serial. Here we revisit the time-varying version of this model with our novel parallel FSP implementation.

**TABLE 3.**
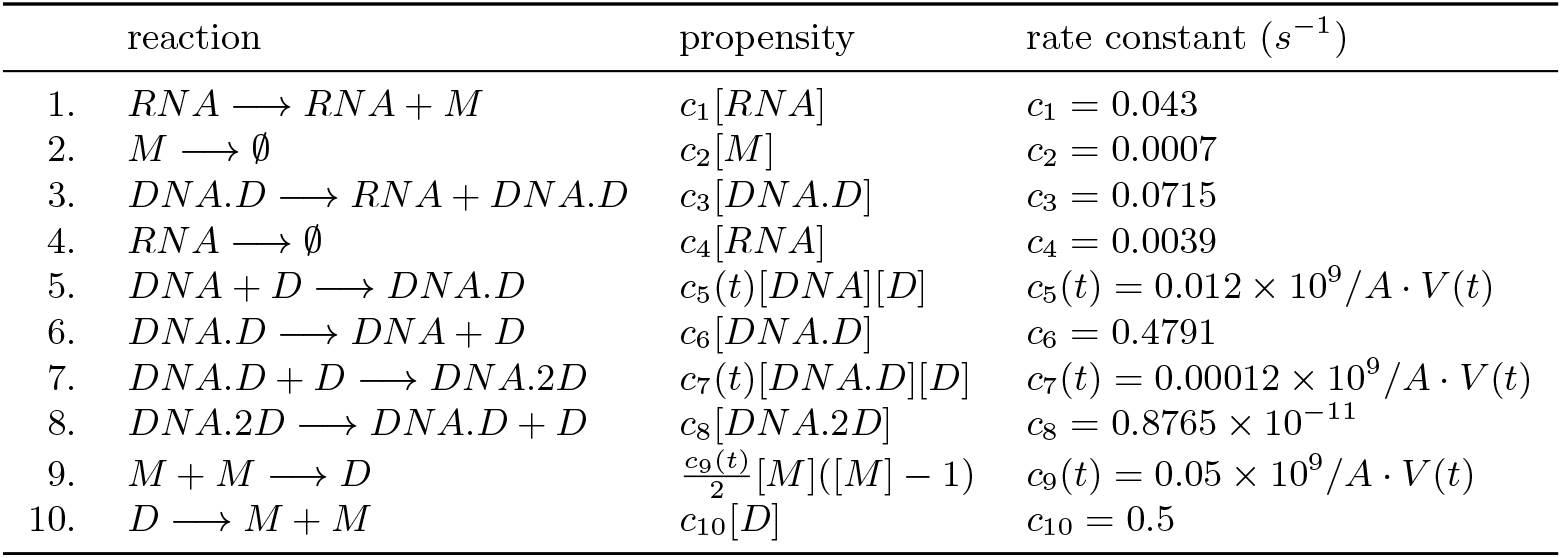
Reaction channels in the six-species transcription regulation model. The parameter A = 6.0221415 × 10^23^ is Avogadro’s number, and V = 10^−15^2^*t/τ*^ L is the system volume chosen for the numerial test. Here, τ is the average cell cycle time, which we set to 35min based on [28] ([X] is the number of copies of the species X.)

| reaction | propensity | rate constant ($s^{-1}$) |
| --- | --- | --- |
| 1. $RNA \rightarrow RNA + M$ | $c_1[RNA]$ | $c_1 = 0.043$ |
| 2. $M \rightarrow \emptyset$ | $c_2[M]$ | $c_2 = 0.0007$ |
| 3. $DNA.D \rightarrow RNA + DNA.D$ | $c_3[DNA.D]$ | $c_3 = 0.0715$ |
| 4. $RNA \rightarrow \emptyset$ | $c_4[RNA]$ | $c_4 = 0.0039$ |
| 5. $DNA + D \rightarrow DNA.D$ | $c_5(t)[DNA][D]$ | $c_5(t) = 0.012 \times 10^9 / A \cdot V(t)$ |
| 6. $DNA.D \rightarrow DNA + D$ | $c_6[DNA.D]$ | $c_6 = 0.4791$ |
| 7. $DNA.D + D \rightarrow DNA.2D$ | $c_7(t)[DNA.D][D]$ | $c_7(t) = 0.00012 \times 10^9 / A \cdot V(t)$ |
| 8. $DNA.2D \rightarrow DNA.D + D$ | $c_8[DNA.2D]$ | $c_8 = 0.8765 \times 10^{-11}$ |
| 9. $M + M \rightarrow D$ | $\frac{c_9(t)}{2}[M]([M] - 1)$ | $c_9(t) = 0.05 \times 10^9 / A \cdot V(t)$ |
| 10. $D \rightarrow M + M$ | $c_{10}[D]$ | $c_{10} = 0.5$ |

The FSP error tolerance is set to 10^−4^. We start at the initial state

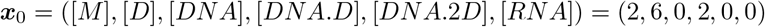

and set the final time to *t*_*f*_ = 5 minutes. The FSP tolerance was set to 10^−4^. For this problem, we use the default hyper-rectangular shape for the FSP-truncated state space. Similar to the previous example, we compare the adaptive and static FSP variants, with the latter using the final bounds output by the former. In addition, we compare the effects of using a simple load-balancing apporach based on distributing states equally, and a graph-based approach. All variants use the BDF method implemented in SUNDIALS [30] for solving the truncated ODEs systems (c.f., eq. (3.7)).

Fig. 8 shows the marginal distributions at the final time, and Fig. 9A shows the size of the dynamic state space. Increasing the number of processes consistently reduce the workload per process (Fig. 9B), and the total workload is distributed almost equally among processors in all variants (Fig. 9C).

**FIG. 8.**
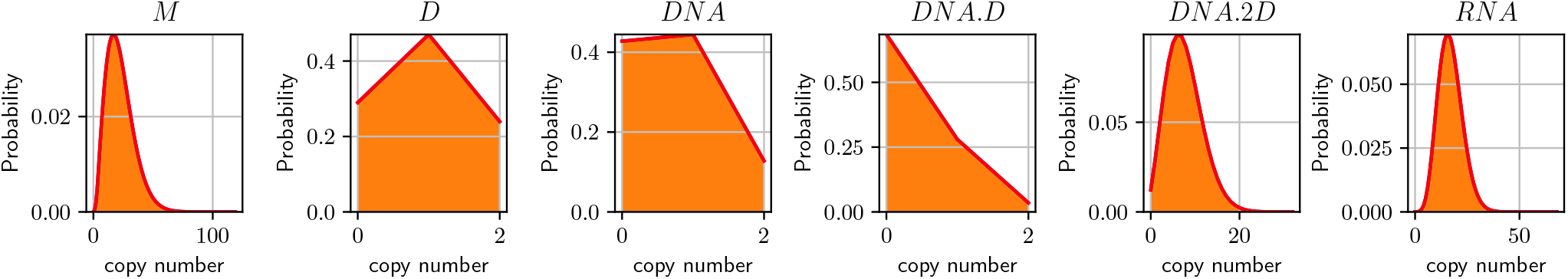
Transcription regulation example. Marginal distributions of monomer, dimer and DNA at time *t*_*f*_ = 5 minutes.

**FIG. 9.**
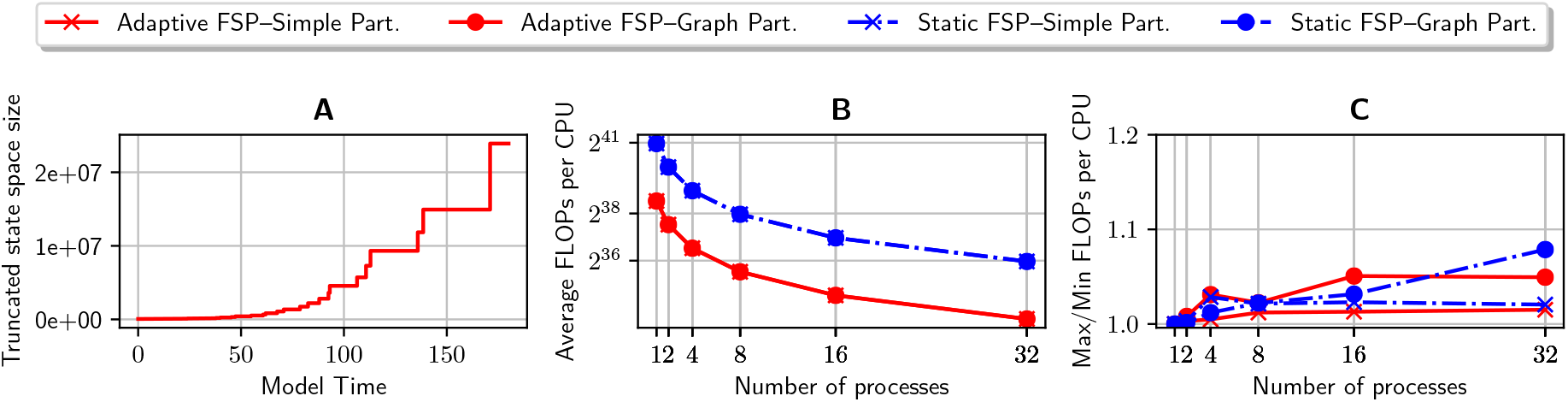
Workload in the numerical integration of the six-species transcription regulation example. (A): Size of the FSP state space over time for the adaptive FSP variants. (B): average number of floating point operations (FLOPs) per process for four different choices of FSP adaptivity and load-balancing. (C): The load-imbalance ratio, defined as the ratio between the maximum and minimum numbers of FLOPs per process, in the four algorithmic variants.

We plot the solver runtime across different number of processors in Fig. 10. Interestingly, using the more advanced graph-based techinique did not improve runtime significantly, with the graph-based partitioning incurring a slight increase in time spent in state expansion (Fig. 11) due to additional overhead for generating the graph. On the other hand, using the adaptive FSP is significantly faster than the static approach (Fig. 11), despite the fact that the adaptive variant has some extra overhead for dynamically re-distributing the states at the intermediate timesteps.

**FIG. 10.**
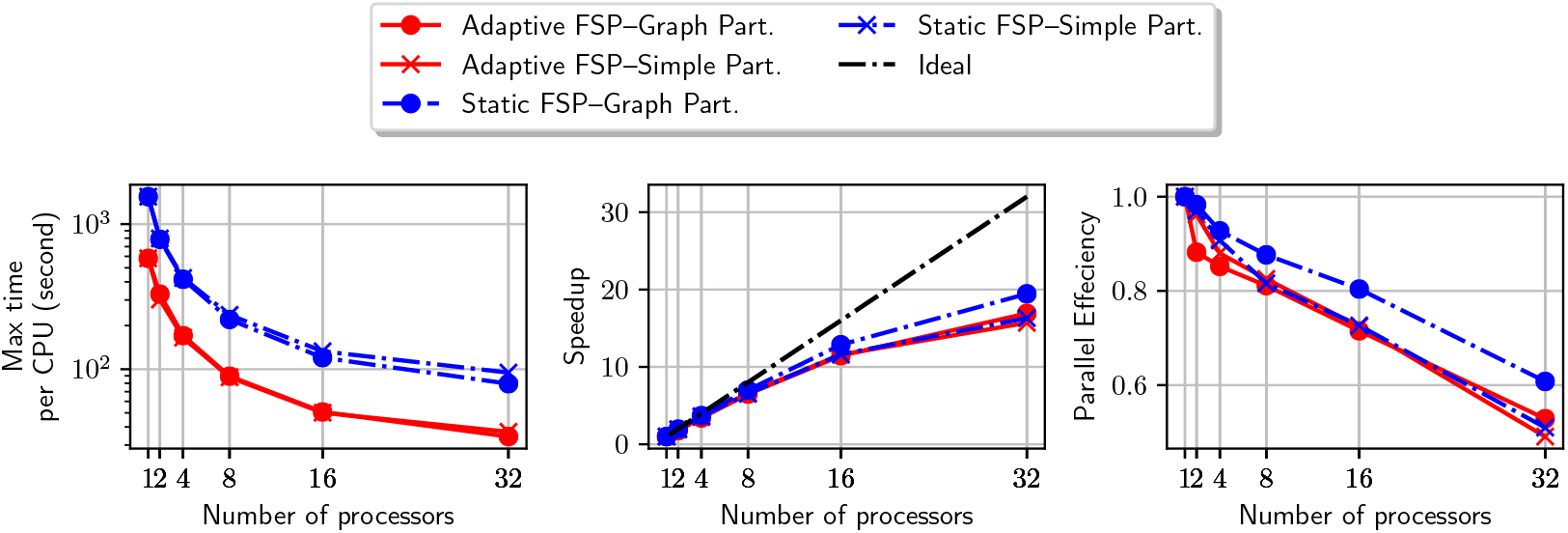
Computational time, speedup, and parallel effeciency in the parallel integration of the transcription regulation example. The four curves correspond to different choices of the FSP adaptivity (either adaptive or fixed), and the load-balancing model (graph-based or simple). The straight diagonal line in the middle plot represents the ideal linear speedup.

**FIG. 11.**
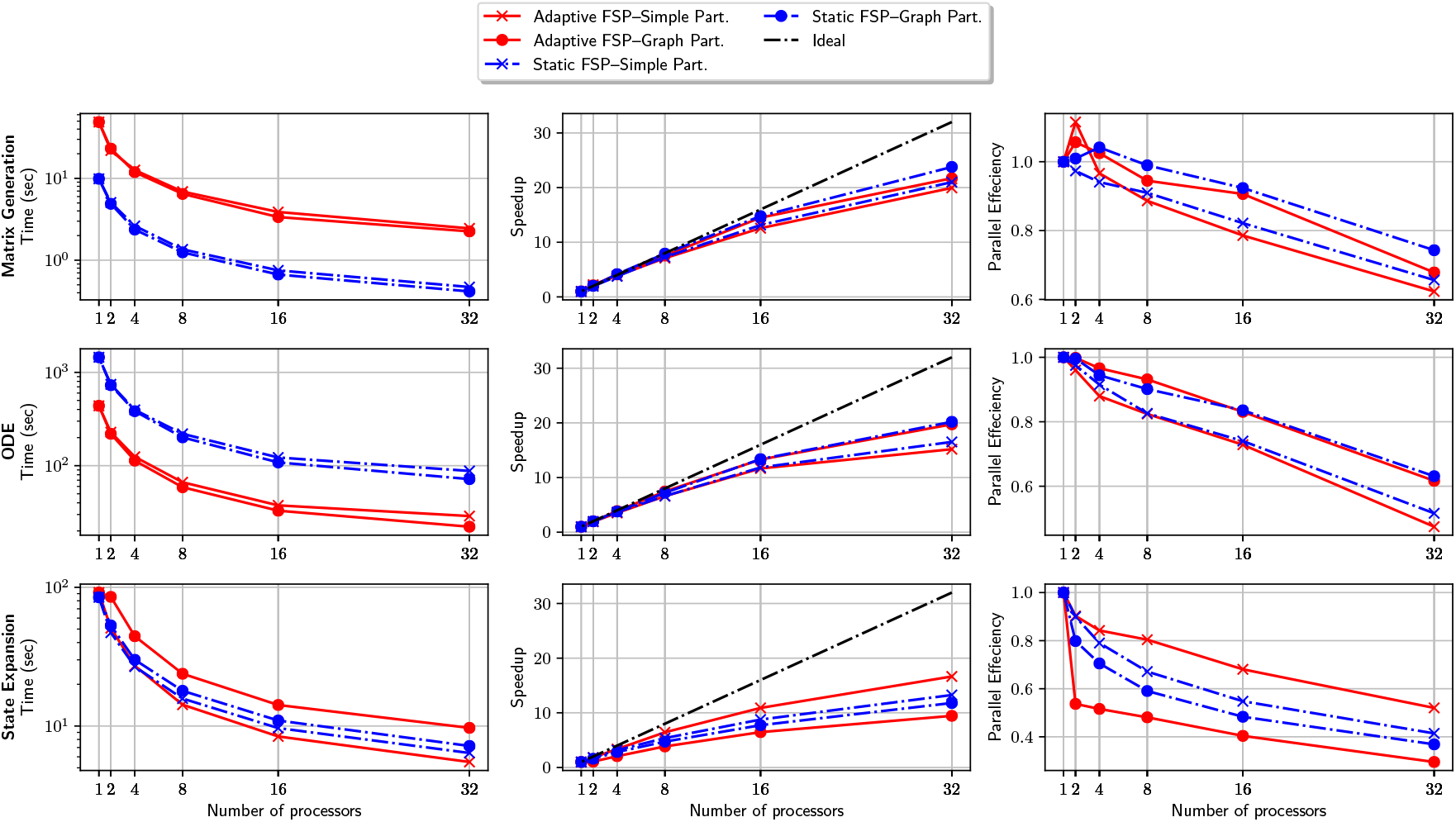
Detail timing for critical components in the parallel solution of transcription regulation example. These critical components include matrix generation (first row), solution of the truncated ODEs systems (second row), and state space expansion (third row). We compare the computational time, speedup, and parallel effeciency of four algorithmic variants based on the choice of FSP algorithm, either adaptive or static, and the choice of load-balancing method, either simple or graph-based.

The single-core performance of our adaptive FSP implementation is comparable, if not faster, than the previous approach reported in [15]. In particular, the experiment reported in [15] solve the same problem in the order of 4 × 10^4^ second, running on a single Intel Xeon E5-4640 CPU at 2.4GHz, while our single-core variants finish the problem in the order of 10^3^ second. This suggests that our FSP implementation could be as effecient as comtemporary methods when running on a single core, and the speedup from using more cores, a unique advantage of our parallel implementation, can significantly reduce the time spent in the forward solution of the CME on this example.

### 5.3. A five-species signal-activated gene expression model

The final model that we consider is an extension of the spatial signal-activated gene expression model that was previously fit to MAPK-activated gene expression data in yeast [43, 45]. We consider a single gene with four states that can transcribe two different RNA species. This could occur, for example, due to overlapping or competing promoter sites. Each RNA species is transcribed in the nucleus, then transported to the cytoplasm where they can be degraded (table 4). Genes only transcribe RNA when in active states (indexed by 2, 3, 4 in the model). The rate of gene switching to inactive state is modulated by the time-varying Hog1p signal, modeled by

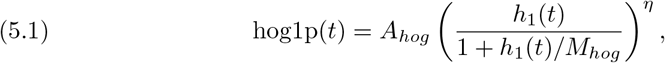

where *h*_1_(*t*) = (1.0 − exp(−*r*_1_ ∗ *t*)) ∗ exp(−*r*_2_ ∗ *t*) and the remaining parameters are *r*_1_ = 6.9*e* − 5, *r*_2_ = 7.1*e* − 3, *η* = 3.1, *A*_*hog*_ = 9.3 × 10^9^, *M*_*hog*_ 6.4 × 10^−4^.

**Table 4.**
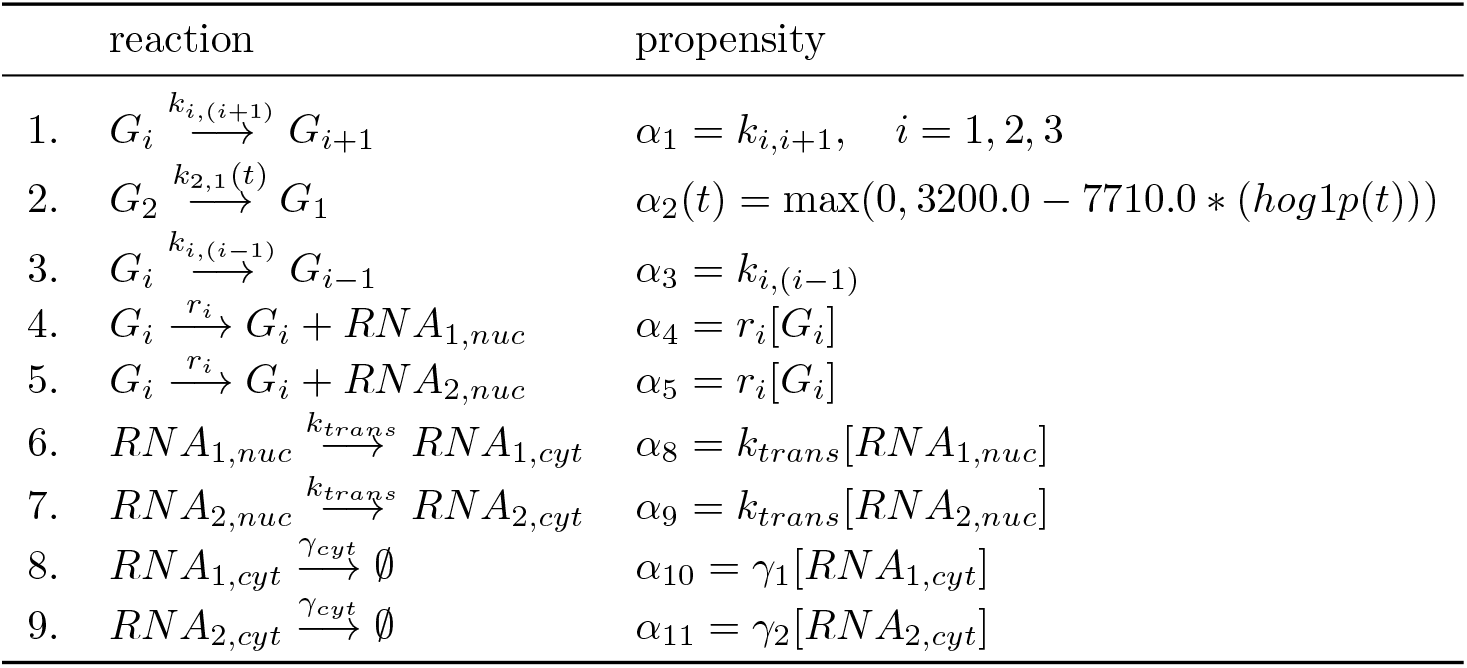
Five-species signal-activated gene expression reactions and propensities. The gene is considered as one species with 4 different states *G*_*i*_, *i* = 0, …, 3.

**TABLE 5.** Parameters in the five-species signal-activated gene expression example. We assume that the time unit is seconds (sec). Hence, the parameters’ units are sec^−1^.

| Parameter | Value |
| --- | --- |
| $k_{12}$ | 1.29 |
| $k_{23}$ | 0.0067 |
| $k_{34}$ | 0.133 |
| $k_{32}$ | 0.027 |
| $k_{43}$ | 0.0381 |
| $r_1$ | 0 |
| $r_2$ | 0.005 |
| $r_3$ | 0.45 |
| $r_4$ | 0.025 |
| $k_{trans}$ | 0.01 |
| $\gamma_1$ | 0.001 |
| $\gamma_2$ | 0.0049 |

We start at

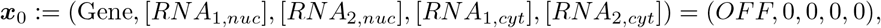

and integrate the CME up to time *t*_*f*_ = 180 second. We use the FSP tolerance of 10^−4^ and an adaptive hyper-rectangular FSP. All variants use the BDF method implemented in SUNDIALS [30] for solving the truncated ODEs systems (c.f., eq. (3.7)).

We compare the runtime of two variants: one using the simple partitioning scheme and the other the Graph-based partitioning scheme. Interestingly, we found that using simple partitioning results in shorter overall runtime across different number of processors, with nearly ideal parallel speedup (Fig. 14). In order to gain more insights into the performance difference between the two variants, we compare the time each variant spends in the two major components of the solver: the ODE integrator time (which includes the cost of matrix-vector multplications), and the state set computation (including state exploration and partitioning). We see that distributing the states using the Graph partitioner indeed result in shorter time for the ODE integrator, as the Graph partitioner seeks to reduce the communication cost of the matrix-vector multplications (Fig. 15). However, this saving in the ODE component is outweighted by the higher cost of the more elaborate partitioning method, which inflates the time spent in the state set component (Fig. 15).

Another interesting observation is that the state set component achieves superlinear speedup when using the simple partitioning scheme. This could be explained by the fact that the number of states needed for the FSP in this example is very large, eventually reaching 23, 908, 221 states (Fig. 14), with the marginal distributions of the RNA having long tails (Fig. 12). This means that the speed of exploring these states with a single core will be limited by memory bandwidth, whereas distributing the exploration among many cores allow for better use of computer memory.

**FIG. 12.**
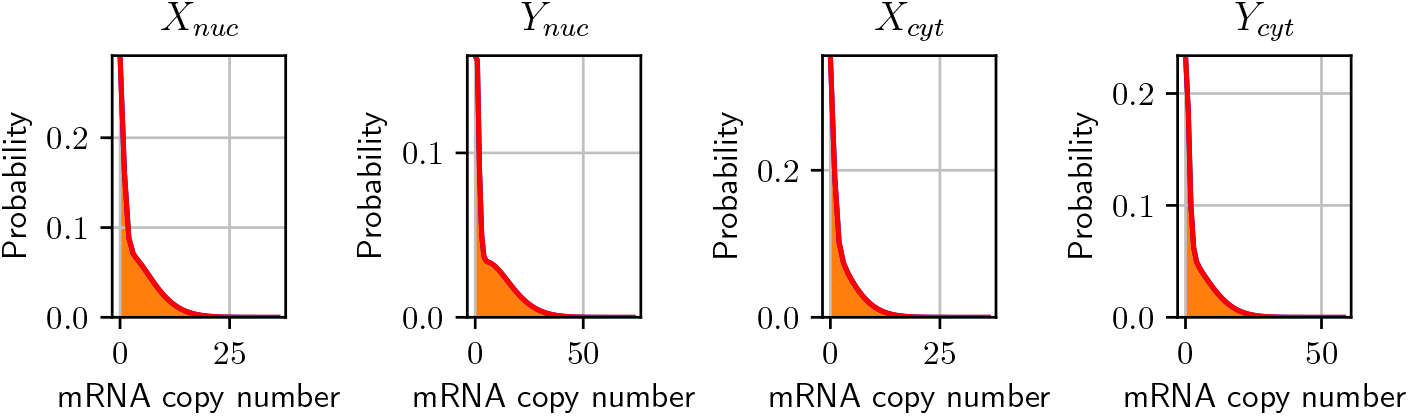
Signal-activated gene expression example. Marginal distribution of the RNA species at 3 minutes after signal activation.

**FIG. 13.**
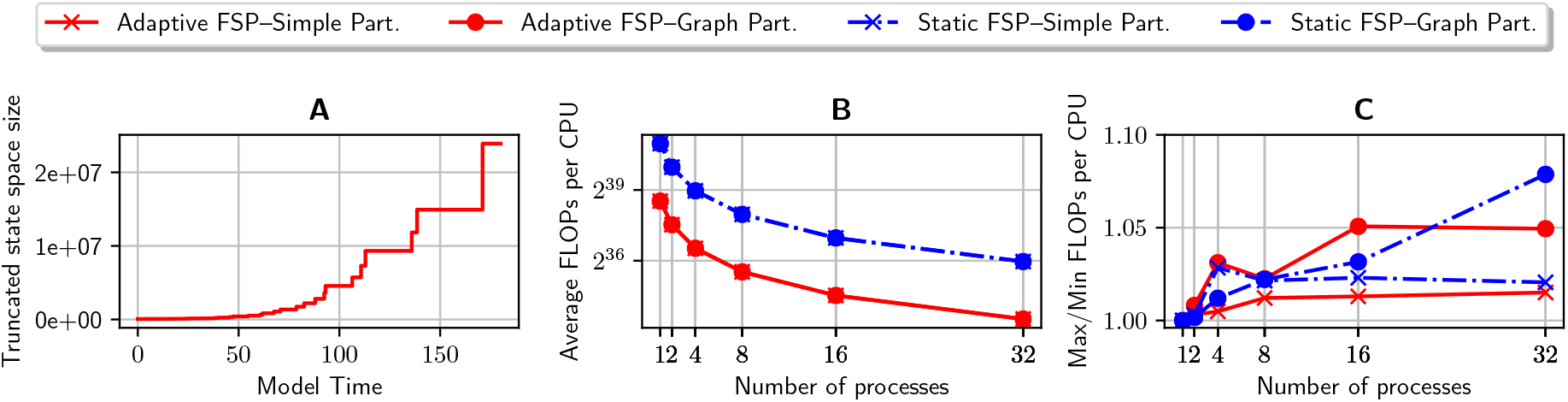
Computational time, speedup, and parallel effeciency in the parallel integration of the signal-activated gene expression example. The four curves correspond to different choices of the FSP adaptivity (either adaptive or fixed), and the load-balancing model (graph-based or simple). The straight diagonal line in the middle plot represents the ideal linear speedup.

**FIG. 14.**
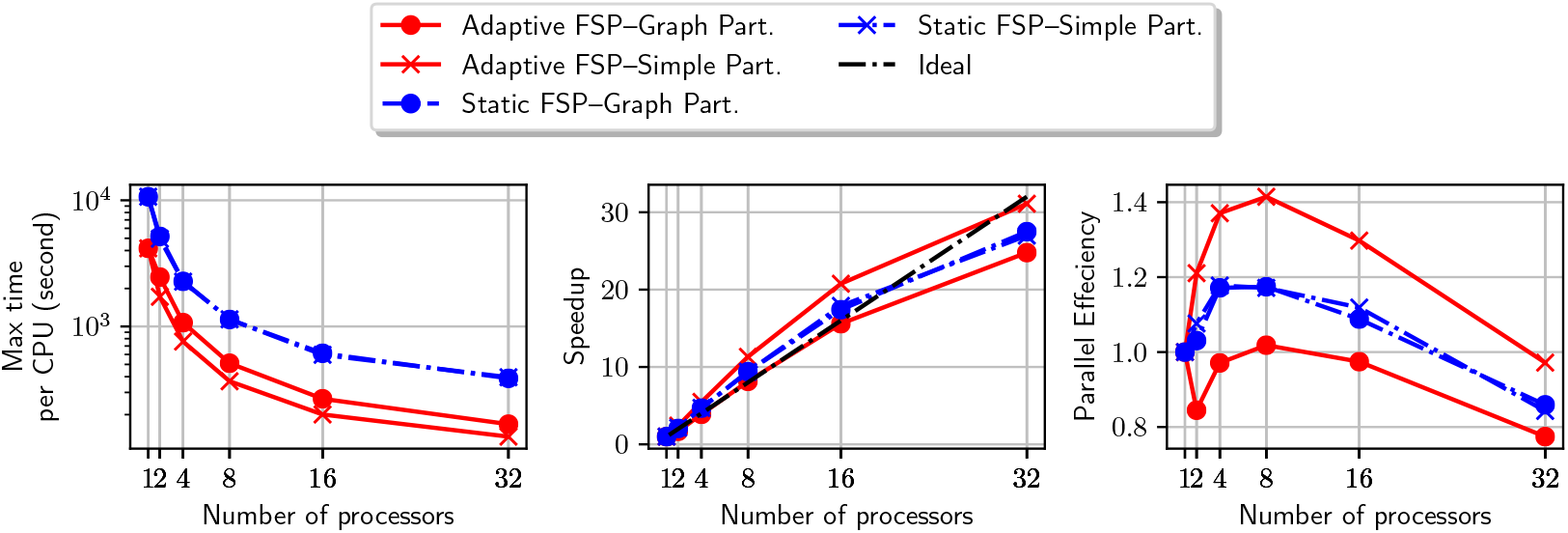
Computational time, speedup, and parallel effeciency in the parallel integration of the signal-activated gene expression example. The four curves correspond to different choices of the FSP adaptivity (either adaptive or fixed), and the load-balancing model (graph-based or simple). The straight diagonal line in the middle plot represents the ideal linear speedup.

**FIG. 15.**
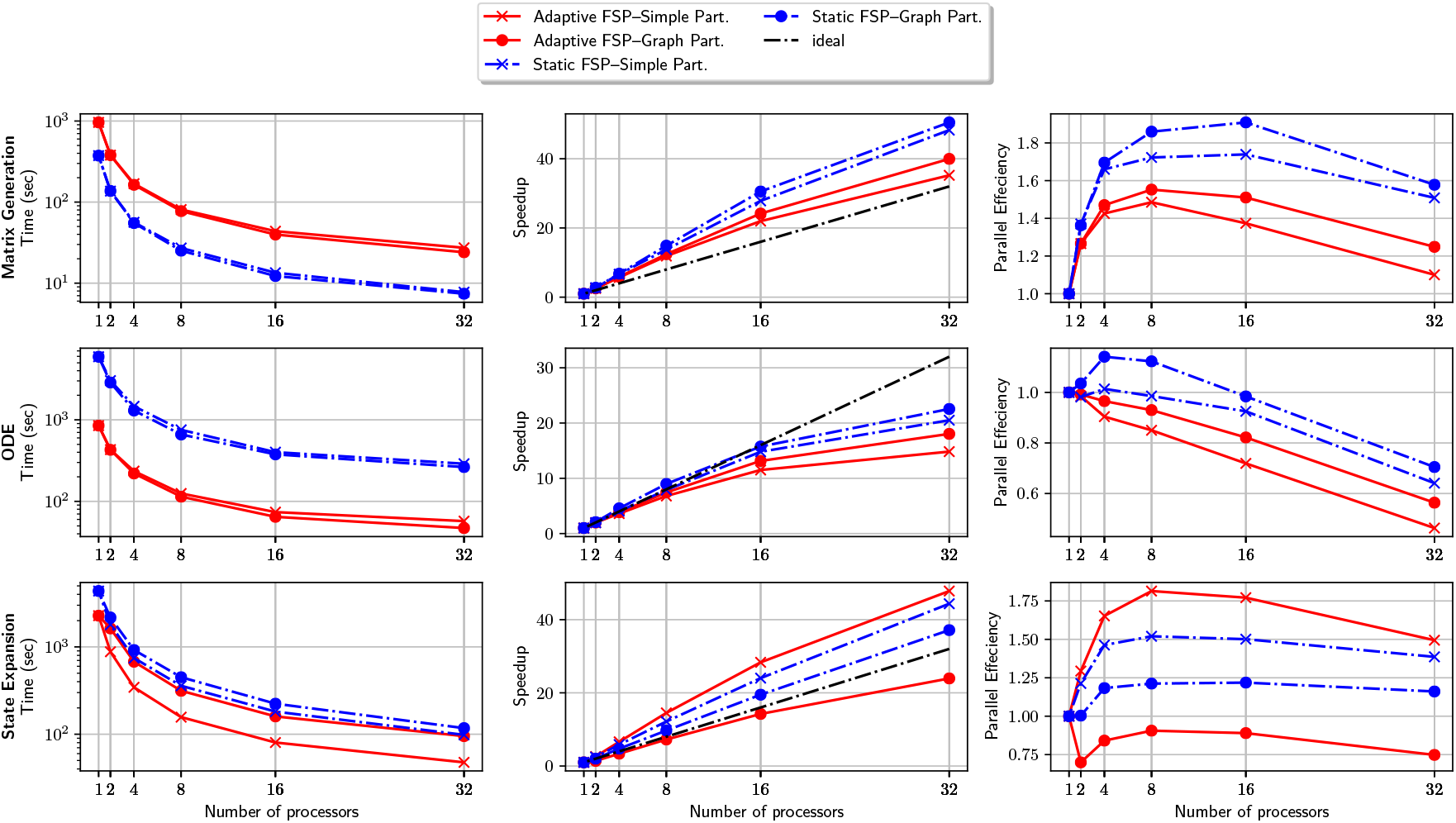
Detail timing for critical components in the parallel solution of signal-activated gene expression example. These critical components include matrix generation (first row), solution of the truncated ODEs systems (second row), and state space expansion (third row). We compare the computational time, speedup, and parallel effeciency of four algorithmic variants based on the choice of FSP algorithm, either adaptive or static, and the choice of load-balancing method, either simple or graph-based.

## 6. Conclusion

In this paper, we presented a novel parallel implementation for an adaptive finite state projection for solving the chemical master equation with time-varying propensities. We test several combinations of FSP state space expansion and load-balancing techniques on sizable problems that require up to 23 million states. Our numerical tests suggest that the largest performance gain comes from the ability of the solver to expand the state space adaptively in parallel, which is a novel feature of our solver in comparison to previous attempts at parallelizing the FSP [52, 58].

While our present implementation has shown significant speedup on realistically large CMEs, there is still room for improvement. First, our object-oriented implementation provides a convenient starting point to parallelize and test alternative strategies for updating the finite state projection, such as sliding window [56] and simulation-driven FSP [51, 54], which have been shown to be competitive with the original FSP in serial settings. Second, recent advances in hybrid MPI-CUDA support in PETSc [40] could provide additional opportunities for the acceleration of the matrix-vector multiplication. Likewise, the task of exploring the state space could utilize the massively parallel GPU cores.

We are exploring the application of our solver to build high performance computational pipelines for the analysis and design of single-cell experiments. For example, we can use our parallel solver to compute more quickly the bounds on the likelihood of single-cell data [21], which facilitates model comparison and selection. It is also possible to make more effecient use of high-performance computing resources for performing Bayesian inference, as we recently explored in [13].

## Acknowledgement

HV and BM were supported by National Institutes of Health (R35 GM124747).

## Footnotes

1 C++ library available at: https://github.com/voduchuy/pacmensl

2 Python wraper available at: https://github.com/voduchuy/pypacmensl

